# Interspecific faecal microbiota transplantation tests microbial contributions to thermal physiology in lizards

**DOI:** 10.64898/2026.10.05.756751

**Authors:** Garazi Martin-Bideguren, Carlos Cabido, Kevin Kohl, Antton Alberdi, Ostaizka Aizpurua

## Abstract

While the influence of microorganisms in the thermal physiology of animals is increasingly recognised, the role of the gut microbiome in enhancing phenotypic plasticity remains underexplored, particularly in ectotherms. Here, we tested the effect of gut microbiomes in modulating the physiological response of lizards to sudden temperature fluctuations. In captivity, we conducted an experiment with wild-caught *Podarcis* lizards: Mediterranean *Podarcis liolepis* (n=9) adapted to warm and dry environments, and Atlantic *P. muralis* (n=18) adapted to cooler and wetter conditions. Animals were assigned to three groups (intervention, control-cold and control-warm) and subjected to microbiota manipulations using antibiotics and faecal microbiota transplants (FMT). In the intervention group, cold-adapted *P. muralis* received faecal suspension from warm-adapted *P. liolepis*, while control-cold and control-warm groups received intraspecific transplants. Microbiome variation was analysed through genome-resolved metagenomics of 162 faecal samples, yielding 313 MAGs across 15 phyla, containing almost 1M annotated genes. One week after transplantation, microbiomes of control animals differed from their original state, whereas those of the intervention group (*P. muralis*), transitioned to a state resembling *P. liolepis*. After two weeks, both control groups showed trends of reversing their microbiomes to their original states. However, the intervention group displayed a distinct trajectory, resembling the microbial donors’. Although potentially relevant trends were observed, these microbiome alterations did not translate into detectable significant differences in their thermal physiology. Nevertheless, our results provide proof of concept for microbial transplants in lizards, and opportunities to further explore the link between gut microbes and host physiological responses in ectotherms.

**Importance:** This study establishes foundational methods for interspecific faecal microbiota transplantation (FMT) in wild reptiles using wall lizards. While microbiota manipulation through antibiotics and FMT has been widely applied in mammals, its feasibility in free-ranging ectothermic vertebrates remains largely unexplored. Although we did not detect significant changes in thermal physiology following FMT, we demonstrate that controlled microbiota manipulation in wild lizards is achievable. This methodological advance opens new opportunities to experimentally test host-microbiota interactions in non-model organisms. Such approaches can facilitate the discovery of previously uncharacterized gut bacterial species and help elucidate how microbial communities contribute to host physiological responses and environmental adaptation. By extending experimental microbiome research to wild reptile systems, this work broadens the taxonomic and ecological scope of host-microbe studies.

## Introduction

Ectothermic heterotherms that undergo large body temperature fluctuations following exposure to environmental heat sources, such as reptiles and amphibians, exhibit complex relationships between ambient temperature and metabolic regulation (1, 2). In arid zones, where there are prolonged periods of heat and drought, reptiles have developed genomic adaptations that confer increased capacities to regulate energy expenditure and maintain metabolic stability (3, 4). However, genomic evolution requires long periods of consistent selective pressures, and therefore not suited for coping with novel sporadic events of thermal stress, which are becoming more common due to climate change (5, 6).

When exposed to sudden environmental changes, ectotherms depend on phenotypic plasticity, expressed through physiological and behavioural adjustments, to cope with adverse conditions (7, 8). Phenotypic plasticity has been historically defined as the capacity of a genotype to express different phenotypes in response to environmental changes (9, 10). However, the recent recognition of the microbiota’s impact on animal host biology has expanded this framework, suggesting that adaptive responses may also be shaped by host-associated microbial communities (11–15). The microbe-driven adaptation hypothesis proposes that gut microbes can help their host adapt to environmental changes more rapidly than the host’s genome alone (13). This hypothesis is supported by evidence that gut microbes influence nutrition, immunity and behaviour (16), and can rapidly modify their functional traits under environmental changes (17), thereby contributing to host phenotypic plasticity (18–20).

While recent studies have reported differences in the gut microbiota of lizards from contrasting temperature environments (21–23), evidence for a causal connection between microbiome differences and animal fitness remains lacking. A systematic review (24) showed that most studies suffer from methodological limitations that impede our understanding of the relationship between microbiomes and animal adaptation. In particular, most studies rely on amplicon sequencing which offers limited capacity to elucidate the microbial functional roles (25, 26) and on observational designs, preventing definitive conclusions about the effects of microorganisms on host biology. To establish causal relationships, experimental manipulation of microbiota is necessary (12), together with measurements of host performance and fitness. Among the few studies implementing manipulative approaches, researchers reported links between gut microbes and thermal tolerance, performance and fitness in tadpoles (27, 28), as well as increased immune capacity in lizards under climate warming (21, 23, 29). However, the mechanisms underlying these effects remain unresolved. Integrating functional metagenomics with microbiota manipulation and physiological tests, researchers can more effectively uncover how microbes enhance vertebrates’ ability to adapt or acclimate to changing environments.

Understanding thermal-microbial relationships is pressing, given ongoing climate-driven declines in reptile populations (30). Here, we studied two *Podarcis* lizard species with contrasting thermal ecologies (31) as a comparative framework for tracking microbiota shifts. We first characterised the community structure and functional potential of the gut microbiome using genome-resolved metagenomics. Next, during a four-week controlled captivity experiment in which we transplanted gut microbiota from warm-adapted lizard donors (*P. liolepis*) into cold-adapted recipients (*P. muralis*) and quantified host metabolic responses via respirometry. In doing so, we aimed to test whether microbes facilitate thermal acclimation and identify potential mechanistic pathways linking microbial functions to lizard thermal tolerance.

## Methods

### Sampling and husbandry

In June 2023, we conducted field trappings in two locations with distinct bioclimatic conditions in the northern Iberian Peninsula, Europe. We captured 18 adult males of *Podarcis muralis*, hereafter referred to as cold-adapted lizards, from a cool, wet Atlantic coastline (Aiako Harria; 18°C, 115 mm precipitation) and nine adult males of *P. liolepis*, hereafter referred to as warm-adapted lizards, from a warmer, drier Mediterranean bioregion (Lizarra; 42.6 N, 2 W; 26°C,30 mm). Lizards were lassoed, individually bagged, and transported to the ZIBA Animal Experimentation Facility (Zarautz, Basque Country). The first faecal sample from each individual was collected to characterise the wild microbiome of each experimental group. Samples were preserved in 1 ml DNA/RNA Shield (Zymo, USA) and stored at −20°C until DNA extraction.

Upon arrival at the facility, lizards were measured, weighed and washed in warm water to remove ectoparasites. Lizards were individually housed in a methacrylate cage [500×150×300 mm] with coconut fibre bedding, a resin rock, two plastic refuges and a 25W lamp to provide a 12/12 hour day/night cycle. Water was offered *ad libitum* and lizards were fed one live cricket (*Acheta domesticus*) every two days.

### Experimental design and sample collection

The experiment lasted four weeks (Fig. 1). Animals belonged to two species: the warm-adapted *Podarcis liolepis* (n=9) and the cold-adapted *P. muralis* (n=18). During the first five days, lizards were acclimated to the experimental facility in two Memmert HPP750Life chambers. *P. muralis* individuals were maintained at 16°C and 80% relative humidity (RH), whereas *P. liolepis* individuals were maintained at 28°C and 40% RH. Following acclimation, animals underwent two days of antibiotic treatment and two days of faecal microbiota transplantation (FMT) (Fig. 1a). All *P. liolepis* individuals were assigned to a warm-control group that received transplants from other warm-adapted animals and remained under warm conditions throughout the entire experiment. Lizards from *P. muralis* were randomly assigned to one of the following groups. The cold-control group (n=9) received transplants from other cold-adapted animals, while the cold-intervention received transplants from warm-adapted animals. Following FMT, both cold-adapted groups transitioned from cool to warm conditions. The temperature and humidity were adjusted gradually at rates of 0.5°C h^−1^ and 1% h^−1^, respectively. Faecal samples were collected and respirometry tests performed at three time-points: at the end of the acclimation, one and two weeks after FMT and environmental change (Fig. 1b).

**Figure 1:**
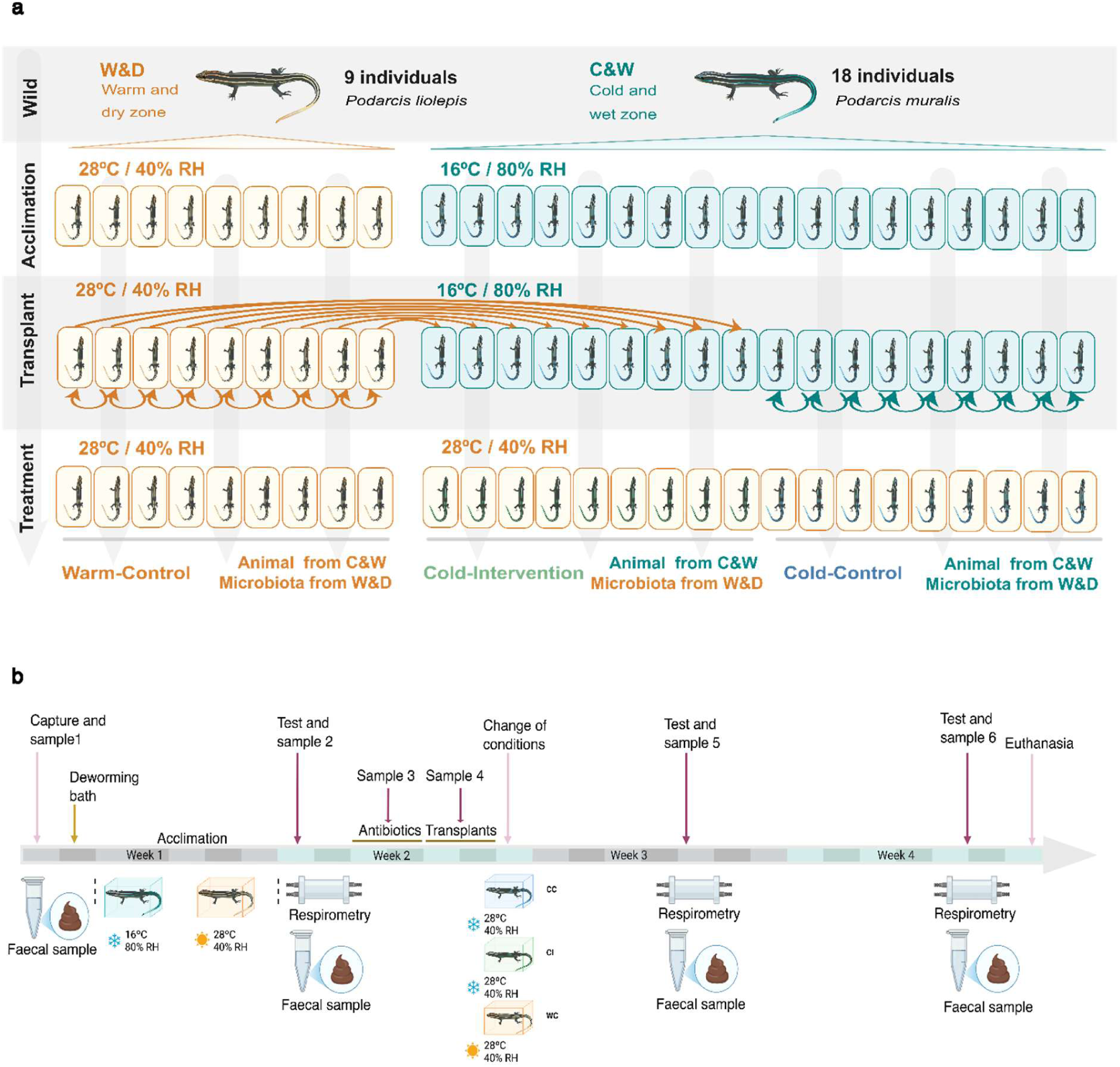
Experimental design and timeline. **a)** The experiment had four phases. First, wild lizards were sampled. Then, acclimated in the animal experimentation facility each one of the species at different conditions. Afterwards, faecal microbiota transplantations were carried out, arrows representing inter- or intraspecies FMT. Finally, during the treatment phase, all lizards were placed at the same conditions. In the wild stage, colours indicate the lizard species: orange (*P. liolepis*) and turquoise (*P. muralis*). Abbreviations correspond to experimental groups: WC (warm-control), CI (cold-intervention) and CC (cold-control). The animal figures have been obtained and modified from the Integration and Application Network, University of Maryland Center for Environmental Science (http://ian.umces.edu/symbols/). **b)** Timeline of the experiment showing the different experimental phases and the time-points at which faecal samples and respirometry measurements were collected. Created with BioRender.

### Faecal microbiota transplantation

Animals were treated for two days with antibiotics prior to faecal transplantation to deplete the native microbiota (Fig. 1a). The antibiotic treatment consisted of a total dose of 75 mg/kg/day that included metronidazole (25 mg/kg/day), neomycin (10 mg/kg/day), ampicillin (20 mg/kg/day), and trimethoprim (20 mg/kg/day). The faecal material for transplantation was collected during the acclimation period and stored in 10% glycerol at 4°C. On the day of the transplant, the faecal material was prepared using previously described method (32). Briefly, the faecal material was homogenised in 300 μl 0.9% NaCl, then filtered and centrifuged. For two consecutive days, each lizard received 20 μl of the faecal suspension (no more than the maximum 2% of their body weight) using a pipette, ensuring that all transplant material was swallowed. The remaining suspension was stored in 1 ml of DNA/RNA Shield (Zymo, USA) for subsequent DNA extraction and sequencing.

### Respirometry

Oxygen consumption rates were measured using closed-chamber respirometry as a proxy for routine metabolic rate (RMR) during 15 minutes. The lizards were confined in a sealed 50 ml tube equipped with internally attached oxygen and temperature sensor spots (OXSP5 and TPSP5, PyroScience) connected via optical fibres (SPFIB-LNS, PyroScience) for contactless oxygen measurement. The fibres were linked to the FireSting-O2 device (FSO2-C4, PyroScience), allowing real-time data collection using the Pyro Workbench software. An empty tube served as a control for measurement-related biases. All measurements were conducted in climate-controlled chambers with lizards in a postabsorptive state and under dark conditions to minimise activity.

### DNA extraction and sequencing library preparation

Tubes containing faecal samples were bead-beaten in a well of the Lysing Matrix E 96-Well Plate (1.2 mL, MP Biomedicals™, cat. number 116984010) for 12 minutes using the TissueLyser. DNA extraction was conducted following the in-house developed DREX protocol (33), using 200 μl of the lysate and obtaining 50 μl of purified DNA (34). Sample processing was conducted in batches of 96 samples along with extraction blanks (water). DNA concentrations were measured using Qubit DNA HS Kit and a Qubit Flex Fluorometer (Thermo-Fisher Scientific). About 200 ng of DNA extracts were sheared using Covaris LE220 machine to around 350 bp-long fragments. Fragmented DNA was then built into sequencing libraries using the custom-developed BEST (35) protocol using Illumina adaptors. Quality of the libraries was assessed using a qPCR assay (Mx3005, Agilent) from where the necessary number of cycles needed for amplification were obtained. Libraries were indexed using dual IDT8 indexes in a 50 μl reaction mix with 10 μl of template DNA. The indexing procedure consisted of 95°C for 12 min, followed by cycles of 95°C 20s, 60°C 30s and 72°C for 40s and finishing up with 72°C for 5 min. Indexed libraries were then purified and quantified using a Fragment Analyzer and pooled equimolarly. Libraries were sequenced using an Illumina NovaSeq X at 150bp paired-end using Novogene (Novogene, UK) services.

### Bioinformatics

Sequencing data processing of the 167 samples was carried out using an in-house developed Snakemake-based pipeline (https://github.com/3d-omics/mg_assembly). First, reads were pre-processed to trim adaptors and low-quality sequences using fastp 0.23.4 (36). After indexing the reference genome of *Podarcis muralis* (PodMur_1.0, NCBI Accession Number GCF_004329235.1), quality-filtered reads were mapped against the host genome using Bowtie2 2.5.1 (37), and unmapped reads retained for subsequent assemblies.

Individual animal-level coassemblies were carried out by merging all the samples from each lizard. Megahit 1.2.9 (38) was used for metagenomic assembly, followed by ensemble binning using CONCOCT 1.1.0 (39), MaxBin2 2.7.7 (40) and MetaBAT2 2.15 (41), and bin refinement using MAGScoT (42). All the refined bins were dereplicated together with dRep 3.4.3 (43) using a 98% average nucleotide identity (ANI) clustering threshold. After CheckM2 (44) and quast 5.2.0 (45) quality assessment, only the metagenome-assembled genomes (MAGs) with completeness over 70% and contamination below 10% were used to build the bacterial genome catalogue employed for downstream analyses. Genome quantification was carried out by mapping the metagenomic reads of each sample to the reference genome catalogue using Bowtie2, followed by quantification using coverM (46). Samples microbial fraction was assessed using SingleM (57, 58) and the Domain-Adjusted Mapping Rate (DAMR) was calculated as the proportion of reads mapping to the MAG catalogue divided by the fraction of the community predicted to be bacterial or archaeal (58).

The MAGs included in the genome catalogue were taxonomically annotated and phylogenetically placed using GTDB-tk 2.3.2 (47). Functional annotation was performed with DRAM 1.4.6 (48), which includes CAZY (49), MEROPS (50), UniProt (51), KEGG (52) and Pfam (53) databases. The raw annotations were then converted into quantitative genome-inferred functional traits (GIFTs) using R package distillR (54). The reference database contained over 300 metabolic pathways and modules derived from KEGG and MetaCyc (55) databases, which were used to obtain quantitative estimates of the metabolic capacities of the MAGs through assessing the relative abundance of genes required to perform a metabolic task. The distilled GIFTs values were corrected for MAG genome completeness to minimise functional biases (56) and then averaged to obtain overall metabolic capacities at genome level or MCI (Metabolic Capacity Index) values.

### Data analysis

All statistical analyses were performed in R v.4.4.1 (59) and compiled into an R Markdown HTML render to ensure full reproducibility. Alpha and beta diversity analyses were performed under the Hill numbers framework (60), using different combinations of orders of diversity (q) and regularity in the hilldiv2 package (61). We estimated richness at *q = 0*, abundance-weighted neutral diversity at *q = 1* and phylogenetic diversity at *q = 1*, incorporating the evolutionary relationships among MAGs.

Alpha diversity of both lizard species at the same time-point was compared using a parametric t-test when data was normally distributed, and variance between groups was equal. When both assumptions were not met, a non-parametric Wilcoxon test was used. To compare microbial trajectories over time across different groups, generalised linear mixed model (GLMM) or linear mixed model (LMM) were performed using *glmer.nb* and *lme* functions from lme4 (62) and nlme (63) packages respectively, including an interaction between group and time-point, with individual ID as a random effect. PERMANOVA comparisons were made using the *adonis2* function from vegan (64) package. Richness, neutral and phylogenetic dissimilarity matrices were used as response variables, and time-point, population and experimental group were included as explanatory variables. When microbial community trajectories were assessed over time and between groups, PERMANOVA comparisons included interactions between the response variables time-point and experimental group and the *strata* option was included to account for nested data, using the individual ID as a factor. These comparisons were visualised by dbRDA using the *capscale* function from vegan. To assess the differences between post-transplantation samples from the same time-point, across groups, pairwise comparisons were performed using the *pairwise.adonis* function from pairwiseAdonis package (65). These comparisons were visualised using NMDS plots created with ggplot2 v.3.5.2. Functional analysis of the microbiome was performed using distillR (54) and *wilcox.test* function from stats R package to display enriched functional traits across time-points or groups.

Oxygen consumption rates were calculated from Pyro Workbench software (PyroScience, Aachen, Germany) and visualised in R using respR 2.3.3 (66). Only measurements with R^2^>0.8 were retained for analysis and mass-specific oxygen consumption data (ml O_2_ g^−1^ seg^−1^) was calculated for each individual and time-point. To test if the oxygen consumption varied between the experimental groups and over time, we built a linear mixed model using individuals as random effects. In order to detect if any changes in oxygen consumption were driven by FMT or environmental change, we compared the measurements of cold-control and cold-intervention and to perceive the influence of those two events together, we analysed cold-intervention and warm-control lizards’ measurements. Individuals without all the measurements were removed to avoid unbalanced data.

## Results

### Genome catalogue

We generated a total of 4.7 billion sequencing reads from 167 faecal samples collected from 27 animals throughout the experiment, with an average of over 29 million reads per sample. After a preliminary data screening, five samples were excluded from downstream analyses due to low sequencing depth (< 1M reads) and incomplete representation of their microbial communities (singleM < 0.2 & DAMR < 0.3). MAG catalogue constituted 313 MAGs with an average completeness of 92.99 ± 7.17% and contamination of 2.15 ± 2.19% and it spanned 15 phyla, dominated by Bacillota A (119 MAGs), Bacteroidota (77) and Pseudomonadota (34). Among all those reconstructed genomes, only 36 were annotated at the species-level, most of them belonging to Pseudomonadota (33%), resulting in a new species discovery rate of over 88%. The functional annotation of the genomes resulted in 1,002,603 predicted genes, out of which 930,700 (92.82%) received KEGG annotations.

### Gut microbiota comparison of warm-adapted *P. liolepis* and cold-adapted *P. muralis* wild lizards

Analysis of gut metagenomes from wild *Podarcis liolepis* (n = 9) and *P. muralis* (n = 18) detected 214 and 187 bacterial MAGs respectively. Of these, 72 MAGs were unique to cold-adapted lizards, 96 were exclusive to warm-adapted lizards, while 115 MAGs were shared between both species. These genomes belonged to the 15 bacterial phyla found in the MAG catalogue. Despite similar alpha diversity, microbial community compositions displayed marked differences between the two species (PERMANOVA: F_richness_ = 6.625, p = 0.001, r^2^ = 0.209; F_neutral_ = 8.822, p = 0.001, r^2^ = 0.260; F_phylogenetic_= 6.899, p = 0.001, r^2^ = 0.216) (Fig. 2a).

**Figure 2:**
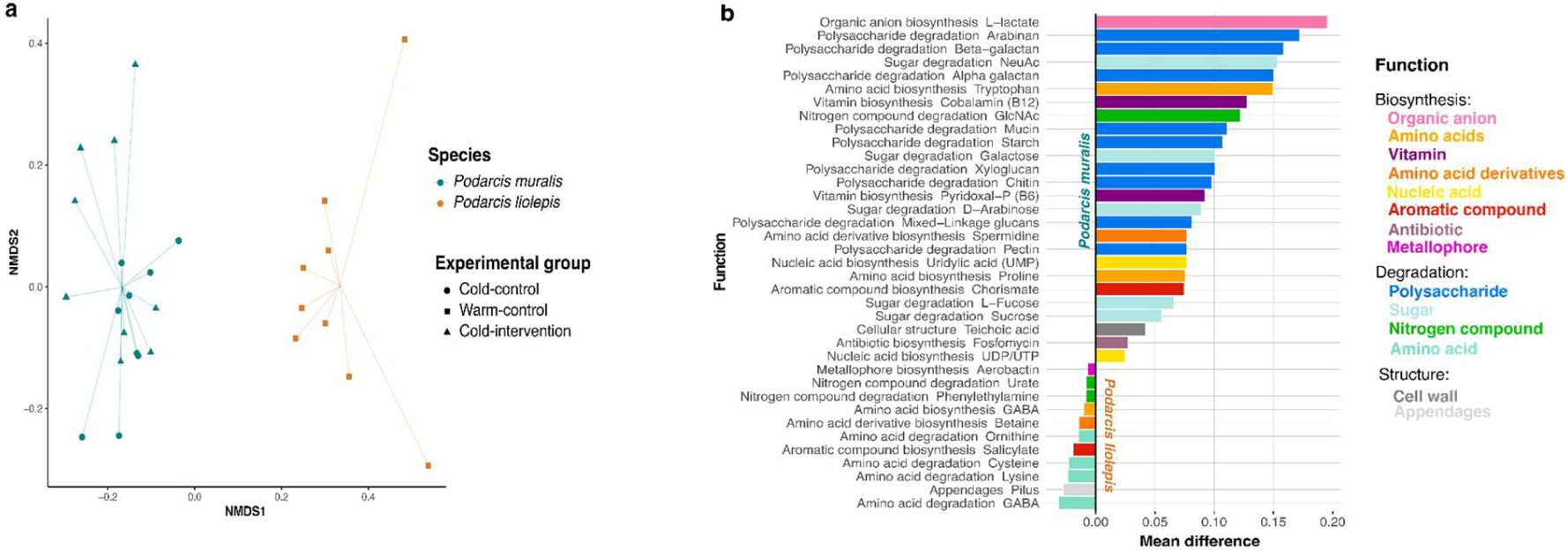
Overview of the community composition and functional capacity of both lizard species in the wild. **a)** NMDS plot of neutral beta diversity, grouped by species and shaped by experiment group. No clear separation among experimental groups is observed in *P. muralis* lizards. **b)** Different microbial functional capacities in each species.

The overall community-weighted Metabolic Capacity Index (MCI), a measure of metabolic capacity at genome level, was significantly higher (t = 2.241, p = 0.034) in cold-adapted lizards (0.346 ± 0.019) than in warm-adapted individuals (0.327 ± 0.024). A total of 37 functional traits showed significant differences between species (Fig. 2b). Cold-adapted lizards displayed an enrichment of 26 functional traits, including degradation of the sugar galactose and the polysaccharide starch and biosynthesis of the amino acid tryptophan. In contrast, warm-adapted lizards showed higher capacities in 11 functional traits, primarily associated with amino acid degradation pathways.

### Effect of captivity on lizards’ microbiota

After acclimation, warm-adapted animals displayed higher diversity than cold-adapted lizards (p < 0.05, Supplementary Fig. S1), although these differences were reduced when phylogenetic relationships among the MAGs were considered (t-test, p = 0.072). On the contrary, captivity did not eliminate the community composition differences observed between species in the wild (PERMANOVA, F_richness_= 5.796, p = 0.001, r^2^ = 0.194; F_neutral_= 8, p = 0.001, r^2^ = 0.25; F_phylogenetic_= 3.867, p = 0.016, r^2^ = 0.138). However, the overall MCI differences observed in the wild were no longer detected after the acclimation period (t = −1.633, p = 0.115). Despite this apparent homogenisation of functional attributes, 16 functional traits remained significantly different between species, including six pathways previously observed in the wild samples (Supplementary Fig. S2), along with an enrichment of GABA and glutamate biosynthesis in warm-adapted lizards.

### Impact of antibiotics on lizards’ microbiota

The lower DNA extraction yields (LMM, p = 0.026, estimate = −1.873) and the higher relative fraction of host DNA in faeces (LMM, p = 4.17e-07, estimate = −0.305) provided robust indirect evidence for a significant reduction in microbial biomass following antibiotic treatment, which aligned with preliminary qPCR analyses that suggested a 60X reduction of biomass. Additionally, we captured a significant decrease in microbial richness and neutral alpha diversity (LMM_richness_, p = 6.02e-10, estimate = −1.152; LMM_neutral_, p = 6e-04, estimate = −11.463), although phylogenetic diversity remained similar to the acclimation phase (LMM_phylo_, p = 0.221, estimate = −0.761). The differences observed in community composition between both species remained after the antibiotic treatment (PERMANOVA, F_richness_ = 3.804, r^2^ = 0.153, p = 0.001; F_neutral_ = 5.541, r^2^ = 0.208, p = 0.001; F_phylogenetic_ = 4.896, r^2^ = 0.189, p = 0.001).

### Microbial changes following faecal microbiota transplantation

To evaluate the potential role of gut microbiota in the thermal acclimation capacity of lizards, we transplanted the gut microbiota of warm-adapted donors (*P. liolepis*) into cold-adapted (*P. muralis*) recipient lizards, *cold-intervention* group (Fig. 1a). Two control groups were established to account for potential effects of the transplantation procedure itself: (1) a *cold-control* group, which received a faecal transplant from another cold-adapted individual, and (2) a *warm-control* group, consisting of warm-adapted lizards that received a transplant from another individual within the same population. After the FMT and environmental change, we tracked temporal changes in gut microbial composition across all experimental groups at one week (FMT1) and two weeks (FMT2) post-transplant.

Alpha diversity varied across the three groups (Fig. 3a). During acclimation, both cold-derived groups showed lower diversity than the warm-control group (p < 0.05). These differences disappeared at FMT1. By FMT2, the cold-intervention group showed a significant increase relative to FMT1 (p < 0.05), surpassing the cold–control group (p > 0.05) and reaching levels comparable to the warm–control group at acclimation (p < 0.05) (Fig. 3a).

**Figure 3:**
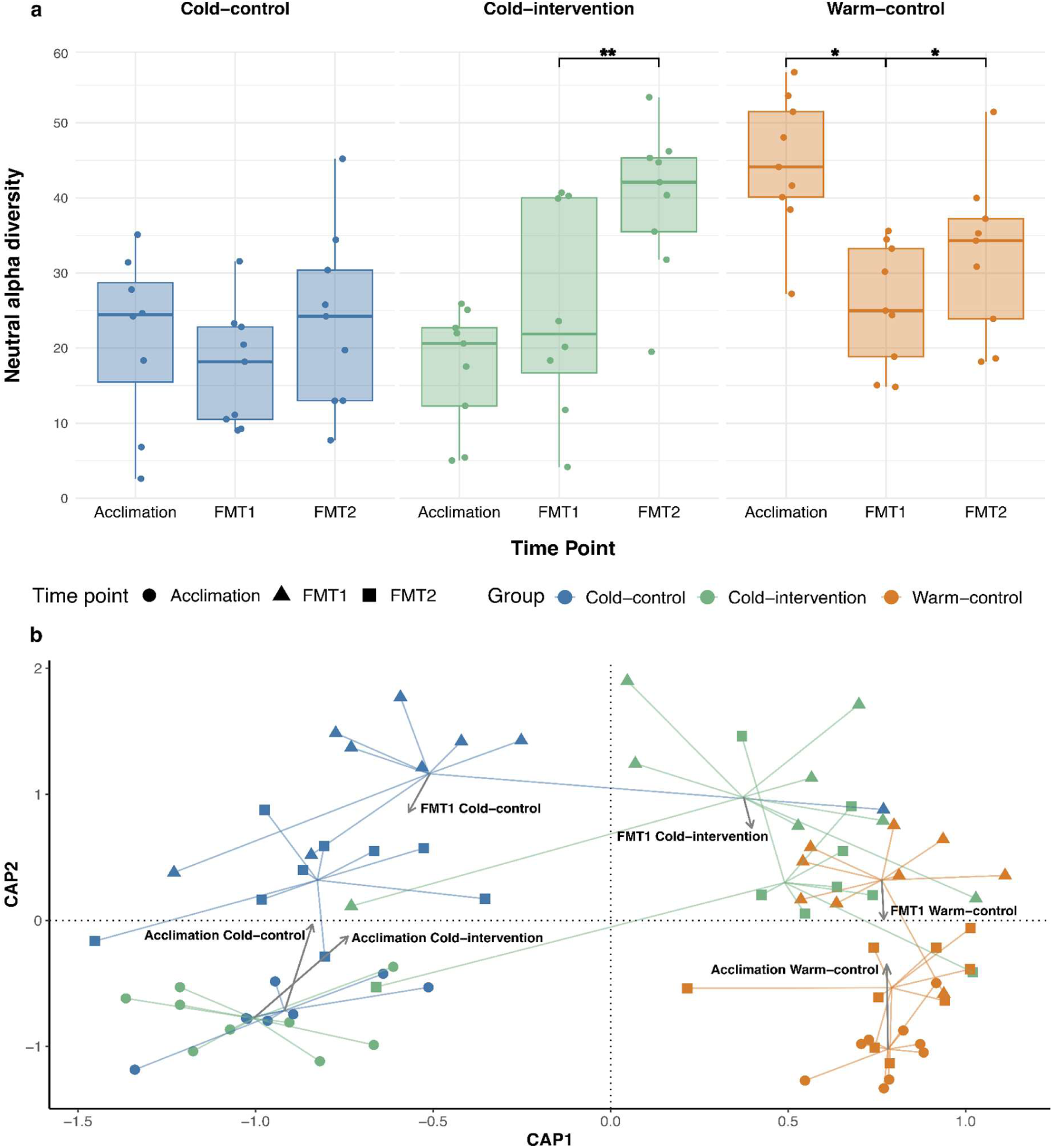
Microbial diversity and community composition variation across three time-points (acclimation, one week postFMT [FMT1] and two weeks postFMT [FMT2]) in the three experimental groups. **a)** Neutral alpha diversity variation across time in the different experimental groups. P-value significance shown by asterisk (*); **= 0.01, *= 0.05. **b)** dbRDA plot of neutral beta diversity for samples collected at the three time-points in the three experimental groups. Arrows indicate the trajectories of the centroids.

The microbial community composition also changed in response to FMT and temperature variation (PERMANOVA, F_neutral_ = 1.861, r^2^ = 0.073, p = 0.001, Fig. 3b). At FMT1, both control groups exhibited only minor deviations in community composition from their acclimation microbiota (PERMANOVA, F_Warm_ = 3.295, r^2^ = 0.17, p = 0.003; F_Cold_ = 3.097, r^2^ = 0.17, p = 0.003) and 11 functional traits were enriched in cold-control (Supplementary Fig. S3). In contrast, the community of cold-intervention group, which received FMT from the warm-adapted (*P. liolepis*) lizards, exhibited more pronounced shifts relative to acclimation (community: PERMANOVA, F = 4.629, r^2^ = 0.23, p = 0.003; two functional traits enriched in FMT1; Supplementary Fig. S4). This group moved away from the cold-control community (p-adj < 0.05) towards that of the donor warm-control community (p-adj > 0.05, Supplementary Table S1) (Fig. 3b). Consistent with this, no significant differences in metabolic capacities (F = 1.371, p = 0.274) or functional traits were detected between the warm–control and cold–intervention groups.

By FMT2, differences among groups became even more pronounced (PERMANOVA, F_richness_= 2.93, r^2^ = 0.19, p = 0.001; F_neutral_= 4.22, r^2^ = 0.26, p = 0.001; F_phylogenetic_= 3.07, r^2^ = 0.20, p = 0.001). Both control groups began reverting toward their acclimation state, whereas the cold-intervention group (*P. muralis*) showed an intermediate state, significantly different from both control groups (p-adj < 0.05, Supplementary Table S2). The comparison of the cold-control and cold-intervention groups revealed two significantly different functional traits. Microbes in the intervention group had a higher capacity to biosynthesise valine amino acid, while the cold-control group showed greater carbapenem antibiotic degradation potential (Supplementary Fig. S5).

### Thermal physiology performance of hosts

Lizard’s physiological performance was assessed by comparing oxygen consumption rates at three time points (Fig. 4). After the acclimation period, the oxygen consumption rates of cold-adapted individuals were significantly higher (Wilcoxon test, p = 0.048; Supplementary Fig. S6) than those of warm-adapted individuals. When analysing the physiological responses to temperature variation in each experimental group separately, we observed that in cold-intervention lizards, oxygen uptake rates decreased significantly at FMT1 (LMM, estimate = −0.0001, p = 0.008, Supplementary Table S3) and FMT2 (LMM, estimate = −0.0001, p = 0.0001, Supplementary Table S4) time-points. In cold-control lizards, however, a significant decrease in oxygen consumption was only detected at the FMT2 time-point (LMM, estimate = −0.0002, p = 0.008, Supplementary Table S4). In contrast, warm-control lizards did not show any change in oxygen consumption rates over time (LMM, p > 0.05, Supplementary Table S5). Nevertheless, when comparing the oxygen consumption rate variation over time across the experimental groups, no significant changes were detected in either of the time-points between the cold-control and cold-intervention groups (p > 0.05, Supplementary Table S6) or between the warm-control and cold-intervention groups (p > 0.05, Supplementary Table S7).

**Figure 4:**
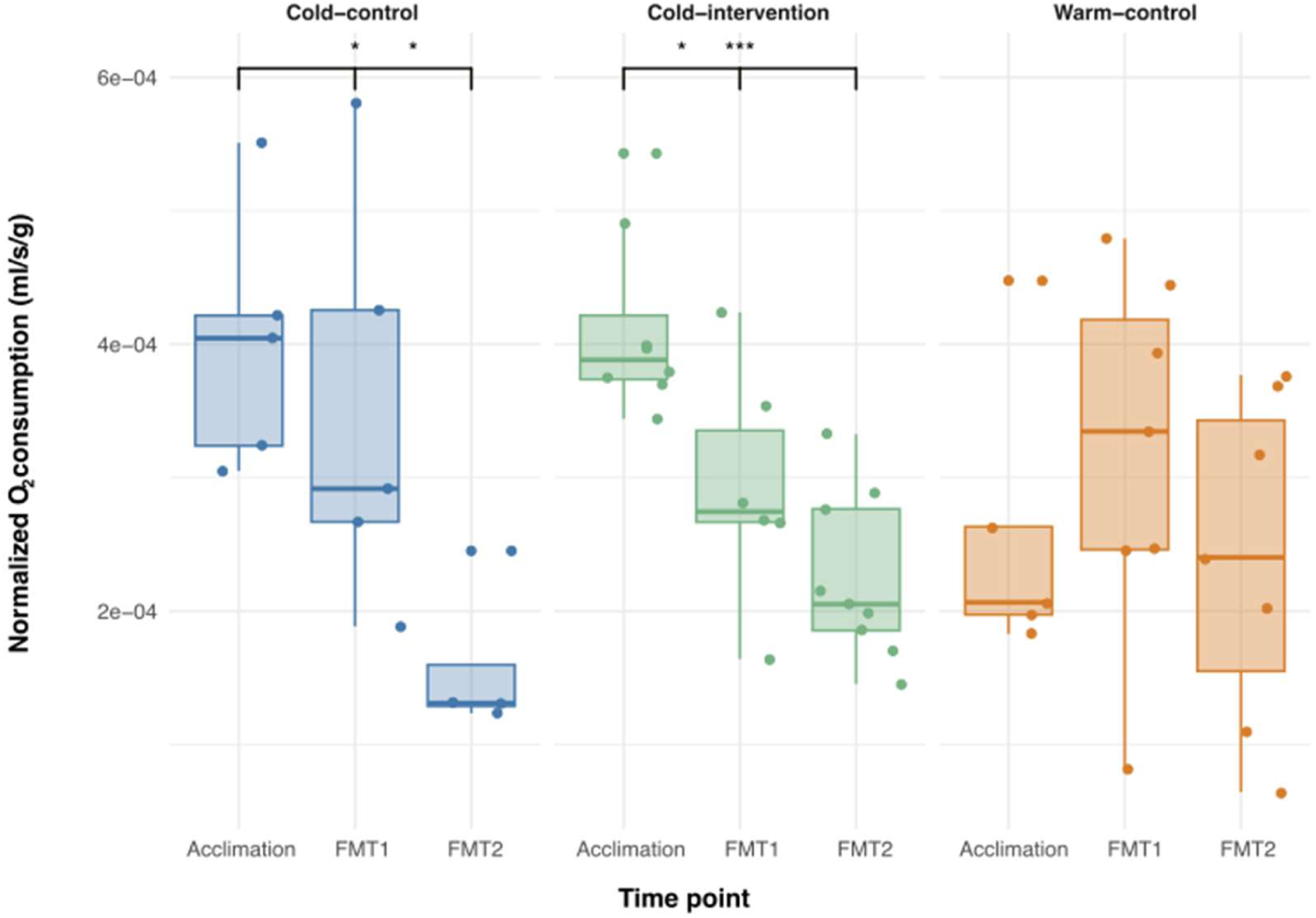
Normalised oxygen consumption rates variation across time in each of the experimental groups. P-value significance shown by asterisk (*); ***= 0.001, *= 0.05.

## Discussion

Experimental approaches for understanding microbial contributions to the thermal physiology of lizards are currently limited, and the causal connection between microbiota variation and host physiology is still largely unexplored, especially in reptiles and other ectotherms. As a significant contribution to that endeavour, our faecal microbiota transplantation experiment in warm- and cold-adapted lizards enabled a robust examination of microbial community shifts and their potential consequences on thermal tolerance. Our results show that, although the transplants successfully altered the gut microbiota, they did not result in a stable community within the studied time period, and they did not exert statistically significant changes in host physiology.

### Genome-resolved metagenomics reveals extensive novel diversity in *Podarcis* gut microbiota

For the first time, we characterised the gut microbiota of wild *Podarcis* from two closely related species with distinct environmental requirements using genome-resolved metagenomics. Unlike 16S rRNA amplicon sequencing, this approach relies on assembling sequencing reads into larger contigs, and binning them into metagenome-assembled genomes (MAGs) (67). By reconstructing near-complete microbial genomes directly from faecal samples, genome-resolved metagenomics not only enables taxonomic assignment but direct annotation of metabolic pathways and functional genes within individual microbial genomes (68).

At high taxonomic levels, our results were consistent with previous descriptions of these species (69, 70) and other reptiles (25, 71) based on 16S rRNA gene sequencing. However, more than 80% of the bacterial genomes recovered from both species lacked species-level annotation, underscoring the vast diversity of uncharacterised bacterial genomes associated with lizards and wild animals in general (72, 73). Such a high proportion of novel genomes highlights the limitations of reference-based approaches in non-model vertebrates, where available databases are still heavily biased toward humans and other model systems (74–76). This result demonstrates the power of genome-resolved metagenomics for improved community resolution and functional inference in non-model vertebrates.

### Contrasting gut microbiota composition and function in two *Podarcis* species adapted to different environments

Although the two species shared over half of their bacterial genomes, their gut microbial communities exhibited significant compositional and functional differences, likely reflecting adaptation to their distinct habitats (77, 78), genetic background (79) or both. The microbiota of cold-adapted *Podarcis muralis* was enriched in genes involved in the degradation of complex polysaccharides such as starch, pectin, and galactans, suggesting an enhanced capacity to process plant-derived substrates and potentially reflecting differences in resource use in colder environments (80). In contrast, the microbiota of the warm-adapted *P. liolepis* was enriched in functions related to amino acid metabolism and nitrogen compound degradation. These pathways may enhance nitrogen recycling and peptide synthesis, contributing to the regulation of host energy balance (81, 82). Among the traits, GABA production stood out as particularly relevant. GABA levels decline under cold and humid stress in mammals (83), and this metabolite plays a key role in the gut–brain axis, influencing gut motility, stress responses, and metabolic regulation (84). More broadly, enhanced potential for amino acid metabolism could contribute to thermal tolerance, as observed in tadpoles (28). These findings suggest that microbial amino acid metabolism may contribute to maintaining physiological performance in warmer environments.

Captivity induced marked alterations in these communities, as previously observed in other vertebrates, where *ex situ* conditions reduce environmental variation and dietary diversity, leading to microbiome homogenisation (85, 86). After the acclimation, the number of microbiome metabolic traits differing between species dropped from 37 to 16, likely reflecting the simplified laboratory environment in which temperature and humidity were the primary variable factors. Nevertheless, key host-associated microbial signatures persisted, particularly pathways linked to amino acid metabolism. These microbiome changes occurred without detectable signs of stress, as individuals maintained stable body weight and normal behaviour throughout the experiment. The persistence of species-specific microbial signatures under identical housing and diet suggests that host-specific factors—such as physiology, immune filtering, immune-mediated selection, or genetic background— exert a dominant influence on microbial assembly, consistent with observation in other reptiles (87). These findings highlight the resilience of host-associated microbiota and the dominant role of the host in maintaining microbiome identity under environmentally controlled conditions.

### Faecal microbiota transplantation modifies community composition and functional capacity of gut microbes

Differences in microbiota composition among populations living under contrasting environments have led to the hypothesis that gut microorganisms can facilitate host adaptation (88, 89). To test this, we transplanted microbiota from warm-adapted *P. liolepis* donors into cold-adapted *P. muralis* recipients. Although faecal microbiota transplant (FMT) has been widely used in mammals and fish (90–92), to our knowledge this study represents only the second application of FMT in lizards (29) and the first between two lacertid species.

Antibiotic pretreatment, commonly used to promote donor microbiota engraftment (93, 94), significantly reduced microbial load and diversity across major lineages, though not completely eliminating the native microbiota, consistent with observations in other species (95–97). Nevertheless, microbial communities remained distinct between species after treatment, indicating that host-specific microbial signatures persisted despite extensive microbiome depletion.

The FMT successfully shifted the gut microbiome of cold-adapted lizards. One week after transplantation, their microbiota was statistically indistinguishable from that of warm-adapted donors, whereas no comparable shift occurred in cold-control animals, indicating that microbial convergence was driven by FMT rather than environmental acclimation alone. Similar donor-recipient convergence has been documented in human studies (98). By the second week, however, the cold-intervention microbiota diverged from the donor community, and adopted an intermediate state between the donor and native communities. Interestingly, predicted functional capacities remained stable, suggesting that compositional differences did not necessarily translate into altered functional potential. This pattern may reflect functional redundancy within the gut microbiota, where taxonomically distinct microbial communities maintain similar metabolic capabilities (17).

Several mechanisms may explain this outcome, including host genetic background, immune-mediated filtering, and early-life host–microbiome interactions that may constrain microbial establishment. As proposed by the “microbiota on a leash” hypothesis (99), hosts regulate their microbial assemblages to maintain stability rather than allowing complete takeover by foreign microbes. In addition, incomplete depletion of the native microbiota may also have allowed residual taxa to compete with donor lineages, generating hybrid communities. Longer-term monitoring is needed to determine whether continued host–microbe interactions promote gradual assimilation of donor taxa or reversion toward native states.

Beyond the biological findings, our study provides methodological advances for microbiome manipulation in reptiles. Pipette-based administration proved to be an effective, minimally invasive delivery method for antibiotics and microbiota suspensions, avoiding the stress and potential injury associated with oral gavage. Moreover, successful donor microbiota engraftment indicates that storage of faecal material in 10% glycerol at 4°C preserved sufficient microbial viability and community integrity for transplantation, although future work should assess potential taxon-specific effects on microbial survival.

### The lizards’ thermal physiology shows an inconclusive response to transplanted microbial communities

To test whether microbiota variation influenced host thermal physiology, we measured oxygen consumption rates before and after transplantation. Initial measurements confirmed the expected differences between populations: cold-adapted lizards exhibited higher oxygen consumption, possibly reflecting increased energetic demands for maintaining physiological function under cooler conditions.

Warm-control lizards kept at constant temperatures maintained stable oxygen consumption over time, indicating that captivity or time alone did not alter metabolic performance. In contrast, cold-control lizards showed a delayed reduction in oxygen consumption after the temperature increase, whereas cold-intervention lizards—those receiving warm-adapted microbiota—exhibited lower oxygen consumption already at the first post-transplantation measurement. This pattern suggests that transplanted microbiota may have facilitated more rapid physiological acclimation to higher temperatures. Nonetheless, group comparisons were not statistically significant, likely due to limited sample size, baseline variability, and the influence of animal activity during respirometry trials.

These methodological limitations may have masked subtle microbiota-mediated effects on metabolism. Furthermore, oxygen consumption provides only a partial indicator of thermal performance, reflecting energy expenditure but not energy acquisition or assimilation (100). While crucial for understanding how species will respond to climate change (101), disentangling this complex relationship remains a significant challenge.

## Conclusion

Faecal microbiota transplants (FMTs) are a crucial tool for establishing causal relationships between gut microbiome variation and host responses. However, the complexity of gut microbiota structure and dynamics presents significant challenges for effectively manipulating these communities. Our analyses showed that FMT can shift the gut microbiota toward a desired state, although this does not ensure reaching a new stable state that persists in time, as the differences between measurements in one and two weeks after transplant demonstrated. More intense antibiotic treatments that further deplete the native microbiota or longer FMT treatments might contribute to more persistent changes.

Although our findings do not provide sufficient evidence to support the microbe-driven adaptation hypothesis, they offer valuable insights into microbial community dynamics following environmental perturbations and FMT, laying the groundwork for future studies that may uncover clearer links between microbiota shifts and host eco-physiological responses. Future work incorporating larger sample sizes, additional transplant events, refined measures of thermal plasticity (e.g. CTmax, SMR) and longer-term monitoring will be essential to clarify these dynamics, test the microbe-driven adaptation hypothesis and evaluate whether microbiome transfer between warm- and cold-adapted animals could serve as a viable conservation tool.

## Acknowledgements

We are grateful to the Herpetological group of Aranzadi for their invaluable knowledge on reptiles to prepare the experiments and perform the captures. As well as to Gipuzkoako Basozainak for locating the lizard populations and the different volunteers that helped with the lizard captures. Finally, we want to thank Dr. Emily Hardison for her assistance with the metabolic data and Prof. Tom Gilbert for supporting the study.

## Supplementary material

Supplemental Tables S1 to S7 and Supplemental Figures S1 to S6.

## Data availability

Raw data and microbial genome sequences of *Podarcis liolepis* and *P. muralis* derived from wild samples were published in the 1st EHI Data Release (73) under Bioproject PRJEB76898, while the data derived from experimental samples were archived samples under Bioproject PRJEB123744. The code to reproduce all analyses is available as a GitHub repository (https://github.com/alberdilab/Podarcis_adaptation_ms), both rendered as a HTML web-book (https://alberdilab.github.io/Podarcis_adaptation_ms/) and archived in Zenodo with doi: 10.5281/zenodo.22954049.

## Ethics approval

Animal captures were approved by the Regional Government of Navarra and Gipuzkoa under licence codes 409E/2023 and 2022FAUNA0000011901, respectively. Animal experimentation was approved by the Regional Government of Gipuzkoa under the licence code PRO-AE-SS-251.

